# Intraductal infection with H5N1 clade 2.3.4.4b influenza virus

**DOI:** 10.1101/2024.11.01.621606

**Authors:** Ericka Kirkpatrick Roubidoux, Victoria Meliopoulos, Brandi Livingston, Pamela H. Brigleb, Stacey Schultz-Cherry

**Affiliations:** Department of Host Microbe Interactions, St. Jude Children’s Research Hospital, Memphis, TN

## Abstract

The ongoing outbreak of highly pathogenic avian influenza (HPAI) H5N1 of the clade 2.3.4.4b in dairy cows has led to numerous questions including how the virus transmits amongst cattle given the limited respiratory infection. One hypothesis is that the virus is spreading through fomites from udder to udder. In these studies, we demonstrate that intraductal inoculation of H5N1 but not H1N1 influenza virus results in infection in mice. This model will be useful to our understanding of the impact of influenza virus on the mammary gland, the potential as a new route of transmission, and can be used to assess if antiviral treatments prevent infections in the mammary gland.

## Text

In March 2024, highly pathogenic avian influenza (HPAI) H5N1 of the clade 2.3.4.4b was detected in dairy cows in Texas and has since been detected in several other U. S. states^1^. Virus has been detected within cow’s milk, indicating that the mammary epithelium may support viral replication^2^. Virus has also been detected on milking machines, leading to a hypothesis that influenza is spreading through fomites from udder to udder instead of the intranasal route^3,4^. There have been studies using cows to better understand mammary infections, however, the cow model is costly and limited^1,5^. We sought to establish a model for intramammary infections of H5N1 and H1N1 influenza virus in mice.

To test the hypothesis that intraductal inoculation is a route of infection, lactating C57Bl/6 mice were infected with 0.5x mean 50% lethal dose (mLD_50_) of A/bovine/Ohio.B24OSU-439/2024 (H5N1) influenza virus into actively lactating nipples (an average of 7 lactating nipples per mouse). The dams and respective pups were monitored daily to assess clinical signs, weight loss, and survival.

After intraductal infection, 3 out of 6 dams exhibited weight loss ranging from 5-30%, with one mouse succumbing to infection (Figure 1A-B). The deceased mouse had high viral titers in mammary glands, lung, and brain, suggesting systemic spread (Figure 1C). Weight loss in the dams may have been associated with loss of appetite from the infection, we also hypothesize that the lactating mice suffered from “milk drop syndrome” where they failed to produce milk. Additionally, the mice that demonstrated the most significant weight loss lost their pups, which did not test positive for virus. Surviving pups in this model did not test positive for virus at 1- or 3-days post infection (dpi), indicating that the model may not be ideal for evaluating transmission of the virus in milk from dams to pups.

**Figure 1:**
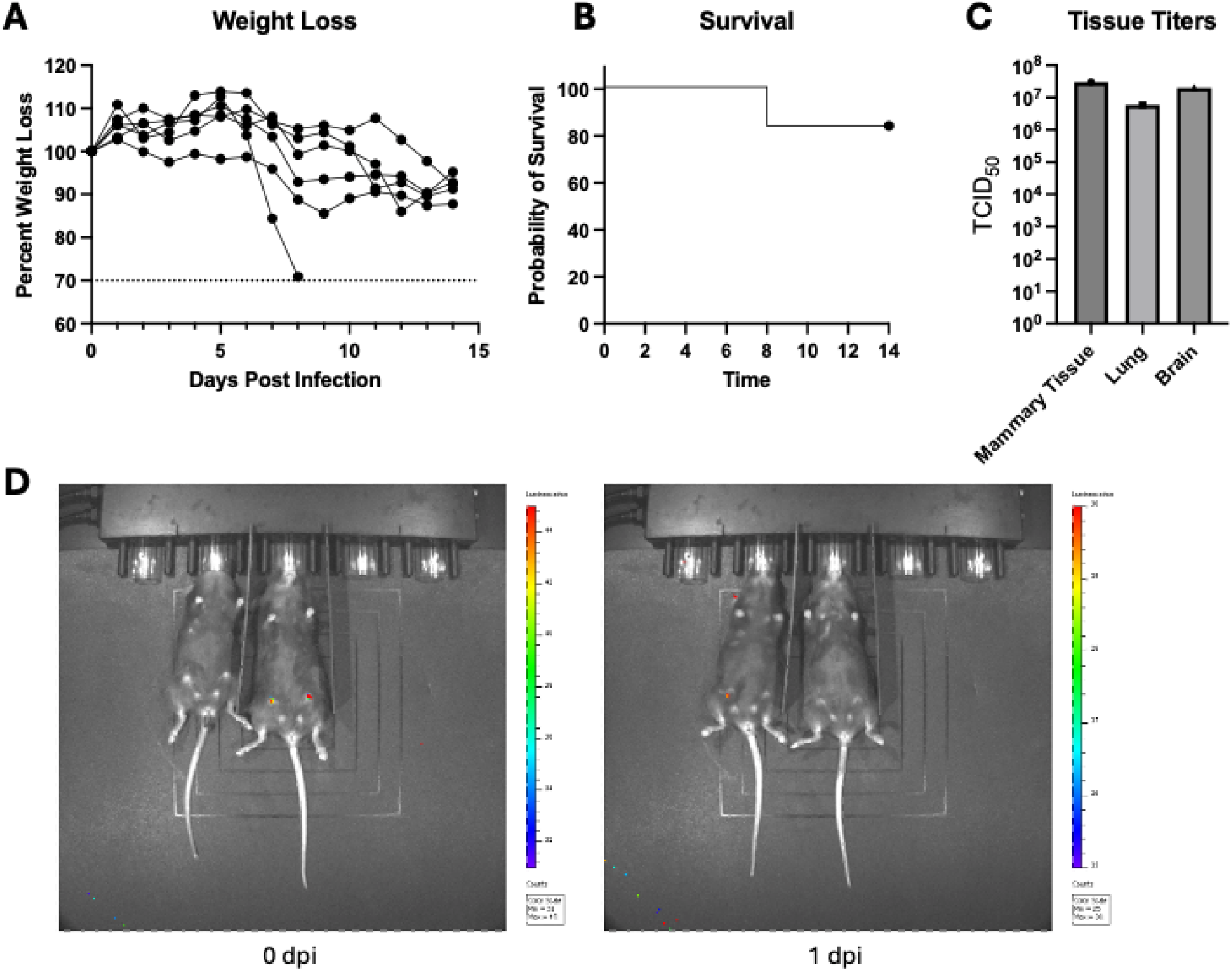
H5N1 and H1N1 intraductal infections. A) Weight loss of mice intraductally infected with H5N1. B) Survival of mice intraductally infected with H5N1. C) Viral titers in various tissues of the mouse that succumbed to H5N1 infection. D) H1N1 intraductal infections. The mouse on the left was uninfected while the mouse on the right received H1N1 via intraductal injection. The image on the left is at 0 dpi while the image on the right is at 1dpi.

To test if mammary tropism is virus subtype specific, we also performed intraductal inoculations in mice using the pandemic H1N1 virus A/California/04/2009. While the injections were successful, the virus was no longer detectable at 1dpi (Figure 1D). These results suggest that H5N1 viruses are more equipped to replicate in mammary glands.

Intraductal inoculations in the mouse model can be used to evaluate lethality and viral spread of H5N1, but not H1N1, viruses. This information is important to our understanding of tissue tropism and can be used to assess if antiviral treatments prevent infections in the mammary gland.

## Acknowledgements

The authors would like to extend their gratitude to Dr. Richard Webby and the Webby lab at St. Jude Children’s Research Hospital for providing the original bovine isolate of HPAI H5N1 (A/bovine/Ohio.B24OSU-439/2024). We would also like to extend our gratitude to the St. Jude Children’s Research Hospital Animal Facility staff and those who operate the Biosafety Level 3 facility. This work was funded by The National Institute of Allergy and Infectious Diseases Contract No. 75N93021C00016 (S.S.-C.), ALSAC (S.S.-C.), National Institute of Health T32AI106700-08 (P.H.B.), and a Ruth L. Kirschstein Postdoctoral Individual National Research Service Award F32AI183804 (P.H.B). The funders had no role in these studies.

## Notes

### Competing Interest Statement

The authors have declared no competing interest.

